# Divergent response to radio-immunotherapy is defined by intrinsic features of the tumor microenvironment

**DOI:** 10.1101/2024.09.06.611679

**Authors:** Jacob Gadwa, Justin Yu, Miles Piper, Michael W. Knitz, Laurel B. Darragh, Nicholas A. Olimpo, Sophia Corbo, Brooke Neupert, Jessica I. Beynor, Alexander Nguyen, Chloe Hodgson, Khalid N.M. Abdelazeem, Diemmy Nguyen, Anthony J. Saviola, Mudita Pincha, Christian Klein, Maria Amann, Sana D. Karam

## Abstract

**Background:** Treatment with immunotherapy can elicit varying responses across cancer types, and the mechanistic underpinnings that contribute to response vs. progression remain poorly understood. However, to date there are few preclinical models that accurately represent these disparate disease scenarios.

**Methods:** Using combinatorial radio-immunotherapy consisting of PD-1 blockade, IL2Rβγ biased signaling, and OX40 agonism we were able to generate preclinical tumor models with conflicting responses, where head and neck squamous cell carcinoma (HNSCC) models responds and pancreatic ductal adenocarcinoma (PDAC) progresses.

**Results:** By modeling these disparate states, we find that regulatory T cells (Tregs) are expanded in PDAC tumors undergoing treatment, constraining tumor reactive CD8 T cell activity. Consequently, the depletion of Tregs restores the therapeutic efficacy of our treatment and abrogates the disparity between models. Moreover, we show that through heterotopic implantations that the site of tumor development defines the response to therapy, as implantation of HNSCC tumors into the pancreas resulted in comparable levels of tumor progression.

**Conclusions:** This work highlights complexity of combining immunotherapies within the tumor microenvironment and further defines the immune and non-immune components of the tumor microenvironment as an intrinsic feature of immune suppression.

**What is already known on this topic:**

- In Head and neck squamous cell carcinomas (HNSCC) and pancreatic ductal adenocarcinoma (PDAC), targeting PD-1 and IL2Rβγ simultaneously (PD1-IL2v) has been shown to be effective when combined with radiation therapy (RT), yet complete response is still limited. The T cell co-stimulatory receptor OX40 (TNFRSF4) has pleiotropic effects, promoting T cell survival, expansion, and memory differentiation in conventional effector T cells, while subsequently limiting regulatory T cell (Treg) suppression by constraining induction and expression of Foxp3. Expression of OX40 is highly upregulated after treatment with PD1-IL2v, and we postulated that combining OX40 agonism with PD1-IL2v and RT would provide additional benefit.

**What this study adds:**

- Using orthotopic models of HNSCC and PDAC, we found that the addition of OX40 agonism unexpectedly drives tumor progression in PDAC, but not HNSCC. Intriguingly, this effect dependent on the tumor microenvironment as the effect is reversed by swapping the location of tumor implantation. This progression was also abrogated by the depletion of regulatory T cells (Tregs), a known mediator of resistance in these models.

**How this study might affect research, practice or policy:**

- Our data demonstrate that unexpected and deleterious effects can stem from combining multiple immunotherapies. These findings hold particular translational relevance as the use of combination immunotherapies is increasingly common on trial.

## Background

Head and neck squamous cell carcinoma (HNSCC) and pancreatic ductal adenocarcinoma (PDAC) are devastating diseases^1–3^, and while many of the underlying characteristics and statistics differ between these two malignancies, the general lack of response to immunotherapy is a unifying feature common to both^4^. While still a boon for some patients, immunotherapy, such as immune checkpoint blockade (ICB), has not unfolded into the magic bullet cancer treatment that was originally predicted. Further confounding our understanding is the notion that ICB and other immunotherapies can elicit drastically different effects across cancer types^5^. Overall, this indicates that tumor-immune interactions in the context of immunotherapy are enormously complex, and that our current understanding remains in its relative infancy.

OX40, encoded by the gene TNFRSF4 and a member of the TNF receptor superfamily, is a secondary costimulatory receptor induced on CD4 and CD8 T cells after T cell receptor (TCR) engagement. OX40 costimulation directly supports the expansion and survival of both CD4 and CD8 T effectors, leading to sustained effector activation, a necessity for durable tumor specific T cell responses^6,7^. While acting as an important activator for effector T cell responses, the overall biological effect of OX40 signaling can be pleiotropic depending on cell type, principal of which being regulatory T cells (Tregs). Tregs have long been established to mediate therapeutic resistance in various cancer types including HNSCC and PDAC, and OX40 signaling has been shown to counteract Treg suppression by constraining the induction of Foxp3, the critical transcriptional regulator for Treg suppressive programming^8,9^. The lack of expression of OX40 on naïve T cells, and the ability to negatively modulate Treg suppression, marks it as an attractive therapeutic target to invigorate tumor reactive immune responses^6,9,10^. Historically however, the success of targeting OX40 has yielded only limited success as a monotherapy, with trials failing to achieve substantial therapeutic efficacy^11^, although there is emerging evidence that OX40 agonism can effectively synergize with ICB^10–13^.

We have previously shown that the immunocytokine PD1-IL2v, which selectively delivers IL-2 signaling via the intermediate affinity IL-2 receptor βγ (IL-2Rβγ) on PD-1 expressing lymphocytes, elicits a powerful antitumor effect in preclinical HNSCC and PDAC tumor models when combined with radiation therapy (RT)^14–17^. Despite significant immune activation, therapeutic response is transient and ultimately incomplete^14,15^. We therefore hypothesized that orthogonal targeting of immune regulatory axes using PD1-IL2v and αOX40 in a combinatorial manner could overcome the transient immune activation we observed.

In this work, we investigate the immunological and therapeutic benefit associated with introducing an OX40 agonist to our radio-immunotherapeutic regimen of RT and PD1-IL2v. We show that OX40 targeting augments the stimulatory effects of RT and PD1-IL2v, further reducing tumor burden in preclinical HNSCC tumor models. In contrast, we report that OX40 targeting acts in opposition in PDAC models and removes the therapeutic efficacy of RT and PD1-IL2v, promoting tumor progression and significantly reducing survival. With this observation came two important findings. First, that accelerated tumor growth of PDAC tumors results from an expansion of OX40^+^ Tregs and diminished CD8 T cell cytotoxicity. Secondly, the disparity in response to combination of RT, PD1-IL2v, and αOX40 is defined by factors specific to the PDAC TME, as heterotopic implantation of head and neck tumors into the pancreas faithfully recapitulates the effect seen in orthotopic pancreatic tumors.

## Methods

### Cell lines

Murine MOC2 and P029 head and neck squamous cell carcinoma and PK5L1940 pancreatic cell lines were used for in vivo studies. All cells were cultured at 37° C and 5% CO_2_ in appropriate media; DMEM-F12:IMDM (1:2) supplemented with 10% FBS and 1% primocin/fungin (InvivoGen), 1.75 ug EGF, 20ug hydrocortisone, and 0.1% insulin for MOC2; DMEM-F12 supplemented with 10% FBS and 1% primocin/fungin for P029; RPMI1640 supplemented with 10% FBS and 1% primocin/fungin for PK5L1940.

### Mouse Tumor Models

Orthotopic HNSCC and PDAC tumor models were established in the buccal mucosa or pancreata, respectively, as previously described^15,18^. Tumors were established at a final concentration of 1×10^5^ /0.1mL per animal for MOC2, 0.5×10^5^/0.1mL per animal for P029, and 2×10^5^/0.02mL per animal for PK5L1940. C57Bl/6 mice and Foxp3^DTR^ mice were obtained through Jackson laboratories. Male and female mice aged 6-10 weeks were used for experiments. Mice were appropriately age and gender matched prior to tumor implantation, and randomized into groups prior to initiation of treatment. For HNSCC models, tumors were measured twice weekly using digital calipers and tumor volume was estimated using the formula (V=A × B^2^/2), where A and B are the long and short diameters of the tumor respectively. All protocols for in vivo animal studies were approved by the Institutional Animal Care and Use Committee (IACUC) of the University of Colorado Anschutz Medical Campus. Mice exhibiting signs of morbidity were euthanized in concordance with IACUC protocol, and primary tumor, serum, and tumor draining lymph nodes were harvested upon sacrifice.

### Antibodies and drugs

PD1-IL2v, FAP-OX40, and aCD25 were provided in collaboration with Roche Pharmaceuticals. PD1-IL2v was injected I.P. at 0.5 mg/kg weekly, FAP-OX40 was injected I.P. at 3 mg/kg at treatment start and again 14 days later, for a total of two doses, aCD25 was injected I.P. at 3 mg/kg weekly. Diphtheria toxin (DT) was administered twice weekly to Foxp3^DTR^ mice at a dose of 1 ug/mouse as previously reported. All antibodies were dosed throughout duration of experiment unless otherwise noted. Treatment was started when average tumor size reached 100mm-150mm^3^ for HNSCC models, or day 7 post implantation for PDAC models.

### Irradiation

Irradiation was performed using the PXi-225Cx image guided irradiator at 225kV, 20mA with a 0.3 mm Cu filter. During mouse buccal irradiation, mice were anesthetized with isoflurane and placed in the prone position. CBCT scans were used for accurate positioning and irradiation was delivered at a dose rate of 5.6 Gy/min. Treatment plans were based on Monte Carlo simulations performed using SmART-ATP software. Buccal irradiation was started when average tumor volume reached 100-150mm^3^. During pancreatic irradiation, mice were placed in the prone position and field beam edges at mouse midline and below ribs. For all in vivo pancreatic experiments, radiation was given at 7 days post implantation.

### Flow Cytometry

Tumors, tumor draining lymph nodes, and blood were collected for flow cytometry. Briefly, tumors were finely minced and placed in Hank’s Balanced Salt Solution (HBSS) containing 200U of Collagenase III and 10ug of DNase I for 30 mins at 37° C, shaking gently every 10 mins. After incubation, tumors were strained first through 70 um filter and then through 40um filter. After centrifugation, red blood cells were lysed using RBC lysis buffer (Invitrogen) and inactivated using HBSS. Lymph nodes were similarly processed into single cell suspension. Blood was immediately centrifuged after collection and resuspended in RBC lysis buffer as described above. Cells were transferred into 24 well plates and stimulated with PMA/ionomycin in the presence of monensin/brefeldin for 4 hours at 37° C. Following incubation, cells were incubated in FC block (CD16/CD32 antibody, Tonbo bioscience) for 20 mins at 4° C, followed by an incubation in a Live/Dead Fixable viability stain kit for 20 mins at 4° C. Cells were then stained with the following antibodies for immune analysis: CD45: PerCP (Clone: 30-F11, Biolegend), CD3: BUV805 (Clone: 17A2, BD Bioscience), CD3: BV786 (Clone: 17A2, Biolegend), CD3: eFluor 450 (Clone: 17A2, Invitrogen), CD4: BUV 496 (Clone: GK1.5, BD Bioscience) CD8: BB515 (Clone: 53-6.7, BD Bioscience), CD44: BV570 (Clone: IM7, Biolegend), NKp46: PE-Cy7 (Clone: 29A1.4, Biolegend), DNAM-1: BV605 (Clone: TX42.1, Biolegend), CD25: BV786 (Clone: C37, BD Bioscience), CD25: BV711 (Clone: PC61, Biolegend), PD-1: BUV615 (Clone: J43, BD Bioscience), PD-1: PerCP-eFluor 710 (Clone: J43, Invitrogen), PD-L1: BV650 (Clone: MIH5, BD Bioscience), OX40: PerCP-eFluor 710 (Clone: OX86, Invitrogen), OX40: BV750 (Clone: OX40, Invitrogen), OX40: Super Bright 436 (Clone: OX86, Invitrogen), Ki-67: APC-eFluor 780 (Clone: SolA15, Invitrogen), Foxp3: Alexa Fluor 532 (Clone: FJK-16s, Invitrogen), IFNγ: BUV737 (Clone: XMG1.2, BD Bioscience), Granzyme B: PE-Dazzle 594 (Clone: QA18A28, Biolegend), IL-10: BV711 (Clone: JES5-16E3, BD Bioscience), GATA-3: Alexa Fluor 647 (Clone: 16E10A23, Biolegend). Samples were analyzed on a Cytek Aurora spectral cytometer (Cytek Biosciences). Data analysis was performed using Flowjo v10. Populations were visualized using FlowSOM and UMAP clustering with Cytobank software (Beckman-Coulter)

### Immune Clustering from Single Cell RNA-Sequencing Datasets

snRNA-Seq data from Hwang et. al. was downloaded from the Broad Institute Single Cell Portal (https://singlecell.broadinstitute.org/single_cell/study/SCP1089/human-treatment-naive-pdac-snuc-seq#/) on April 22nd, 2024. Expression data was imported into R (v.4.4.1) package Seurat (v.5.1.0) , and provided cell type annotations and UMAP coordinates were mapped from the downloaded metadata. Subclustering was performed by subsetting the labeled immune cluster and performing integration on the RNA count data.

scRNA-Seq data from Cillo et. al. was downloaded from the Gene Expression Omnibus under GSE139324 using R package GEOquery on April 30th, 2024. HPV-negative tumor samples were kept for analysis. High quality cells were kept if 200 < nFeature_RNA < 2500, percent mitochondrial reads < 5%, nCount_RNA < 7500, and complexity > 0.75 (defined as complexity = log10(nFeature_RNA) / log10(nCount_RNA)). Integration was performed using Seurat::IntegrateData() with normalization.method = “LogNormalize”.

### RNA-Sequencing

FNA biopsy and post-neoadjuvant surgical patient samples were obtained from the University of Colorado biorepository. All tumors were borderline resectable and treated with neoadjuvant chemotherapy with 30–33.6Gy SBRT, followed by pancreaticoduodenectomy. Tumor samples were reviewed by a pathologist to identify areas of tumor cellularity. Only the pathologist-marked areas of tumor cellularity were scraped and processed per BioSpyder kit instructions. For the human RNA sequencing library preparation, lysates were made from scrapes using BioSpyer FFPE lysis protocol, then diluted in 1:5 in 1X TempO-Seq Lysis Buffer and processed through a TempO-Seq Human Full Transcriptome FFPE Assay 96 sample Kit (BioSpyder, Carlsbad, CA). Reads were aligned and counts generated using Biospyder TempoSeqr platform. Genes with <1 mean raw counts or <1 mean counts per million (CPM) were removed from the data set. Differential expressions were calculated using the limma R package. The resulting fold change was used with the fgsea R package to perform gene set enrichment analysis for Hallmark and C2 Curated gene sets.

### In vitro assays

Healthy donor PBMC were isolated from buffy coats (Zurich Blood Donation Center) using standard density-gradient centrifugation. All primary human samples were cryopreserved. T cell isolation was performed by negative selection using the naive CD4 T cell isolation kit II, human Microbeads (Miltenyi 130-094-131). Naïve CD4 T cell enrichment procedure was performed according to the manufacturer’s instructions. Untouched human CD4 (Miltenyi 130-096-533) and CD8 T cell (Miltenyi 130-096-495) isolation were performed according to manufacturer’s instructions and labeled with 200nM CFSE where indicated. Naïve T cells were labelled with CFSE and cultured on plates coated with anti-human CD3 (end conc. 3µg/ml, eBioscience 16-0037-85) with anti-human CD28 antibody (end conc. 1µg/ml, eBioscience 16-0289-85) at 5×10^5^ cells/ml in AIM-V media (Gibco 12055091). Serial dilution of FAP-OX40 were added - along with huFAP coated beads (2:1 bead to cell ratio) and cytokines rhTGFb (end conc. 2ng/ml , R&D, 240-B-010/CF). Cells were incubated at 37°C, 5% CO2 for 3 days. Cells were incubated with Human TruStain FcX (Biolegend) and fixable viability dye (eBioscience), fixable LIVE/DEAD stain (Molecular Probes by Life Technologies) before flow cytometry. Then, cells were stained in FACS buffer (PBS containing 0.5% BSA) containing flow cytometry detection antibodies for CD4, CD8, CD25, OX40, CCR4, CXCR3, (BioLegend) for 30 minutes at 4°C. Cells were fixed and permeabilized using FoxP3 Transcription Factor Staining Set (eBioscience) and subsequently stained against intra-cellular antigens FoxP3, Bcl-2 (BioLegend) or Ki67 (BD Bioscience) according to manufacturer’s instruction. Samples were acquired with a 5-laser Fortessa (BD) or 5-laser A5 Symphony instrument (BD). Results were analysed with FlowJo v10.6.2. Cell-free supernatants were collected at endpoint and used directly for cytokine measurement stored at -80°C. Cytokines were measured using Cytokine bead assay (BD Biosciences) according to the manufacturer’s guideline.

### Statistical Analysis

All statistical analysis was performed using GraphPad Prism v10. Statistical analyses were completed using one-way ANOVA with Tukey correction for comparisons with 3 or more groups, and unpaired t-test for comparisons with two groups. Kaplain Meier were used to represent survival analysis, using log-ranked Mantel-Cox test for determination of statistical significance. Data is represented as mean SEM, unless otherwise noted. Statistical significance is indicated as *p<0.05, **p<0.005, ***p<0.0005, ****p<0.0001.

## Results

### Tumor infiltrating lymphocytes display increased OX40 expression in preclinical and clinical tumor models post treatment

Previous work by our lab and others have shown that the immunocytokine PD1-IL2v elicits a powerful antitumor effect in various preclinical tumor models^14–17^. However, despite the significant effect on tumor growth, the overall immune response is incomplete and ultimately transient with activation of circulating CD8 T cells diminishing over time (**Supplemental Figure 1A, B**)^14,15^. This prompted us to investigate combinatorial approaches that may augment the tumoricidal potential of PD1-IL2v to increase the overall durability of the response. Examination of flow cytometry data reveals that circulating CD8 T cells have elevated surface expression of OX40 post treatment with RT and PD1-IL2v in preclinical orthotopic models of HNSCC and PDAC (**Figure 1A, B**). Furthermore, tumor infiltrating CD8 T cells have upregulated expression of OX40 in orthotopic models of HNSCC, and Tregs display increased expression in both HNSCC and PDAC models (**Supplemental Figure 1C, D**). Similarly, sequencing analysis from clinical PDAC tumor samples shows that OX40 expression is also increased post irradiation **(Supplemental Figure 1E)**.

**Figure 1:**
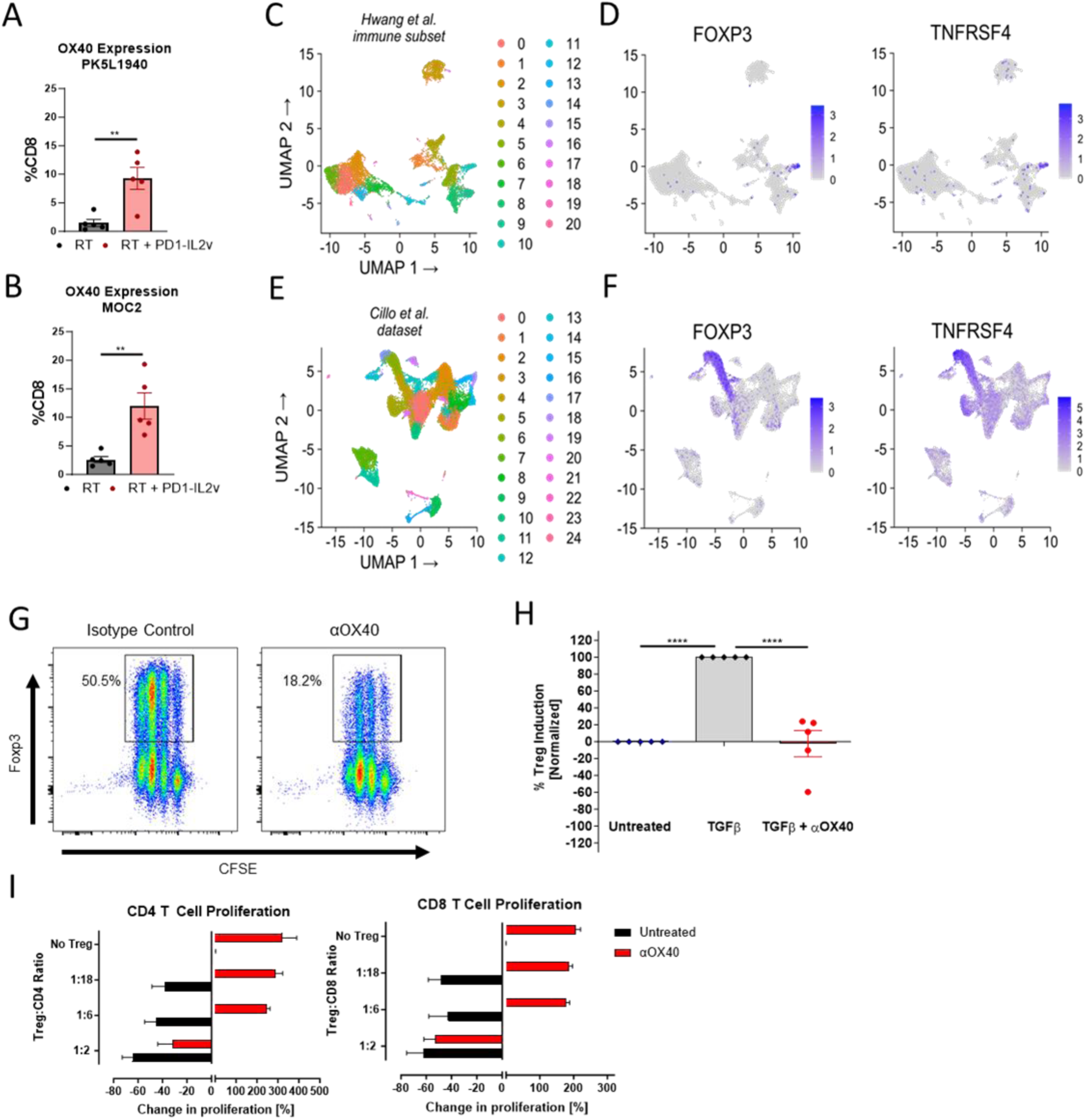
Clinical and preclinical expression of OX40 on CD8 T cells and Tregs in HNSCC and PDAC tumor microenvironments. **A)** OX40 expression on circulating CD8 T cells in preclinical PDAC between mice treated with RT and RT + PD1-IL2v **B)** OX40 expression on circulating CD8 T cells in preclinical HNSCC between mice treated with RT and RT + PD1-IL2v **C)** Uniform manifold approximation and projection (UMAP) analysis of sc-RNA sequencing immune cell clusters from patients with head and neck squamous cell carcinoma at baseline with **D)** with overlaid expression of Foxp3 and TNFRSF4. **E)** UMAP analysis of sn-RNA sequencing immune cell clusters from patients with pancreatic ductal adenocarcinoma with **F)** overlaid expression of Foxp3 and TNFRSF4. **G)** Percent Foxp3 expression in naïve human CD4 T cells treated with 2ng/mL TGFβ and either isotype control or αOX40. **H)** Change in Treg induction in naïve human CD4 T cells cultured with either TGFβ, TGFβ + αOX40, or no treatment. **I)** Percent chance in CD4 and **J)** CD8 T cell proliferation after coculture with Tregs. Naïve CD4 and CD8 T cells were cultured with Tregs at a Treg/effector ratios of 1:2, 1:6, 1:18, or no Tregs and were either untreated or treated with αOX40.

### Tumor infiltrating lymphocytes express OX40 in clinical tumor samples

OX40 costimulation on Tregs has been previously shown to inhibit the induction of Foxp3, the central transcriptional driver of Treg suppression^19,20^. Examining the distribution of TNFRSF4 on tumor infiltrating immune populations in HNSCC^21^ at baseline shows TNFRSF4 expression to be heavily concentrated on Tregs (**Figure 1C, D**). Similar coexpression is observed in immune clustering from PDAC^22^ (**Figure 1E, F**). Of note, expression of TNFRSF4 does not co-colocalize with CD8a in either patient dataset, indicating there is limited CD8 T cell activation at baseline in these patients (**Supplemental Figure 1F, G)**. Based on these data and established literature, we sought to investigate the potential of OX40 as a therapeutic target in combination with RT and PD1-IL2v in preclinical models of HNSCC and PDAC.

### OX40 agonism modulates activation and expansion of T cells in vitro

We next sought to better understand the effects of OX40 agonism on T cell activation and function. To assess the contribution of OX40 signaling in constraining Foxp3 induction naïve T cells labeled with CFSE and cultured with TGFβ in the presence of αOX40 antibody displayed significantly reduced expression of Foxp3 compared to isotype control (**Figure 1G, H**). Furthermore, flow cytometric analysis shows that αOX40 reduces the induction of Tregs when exposed to TGFβ *in vitro* in a dose dependent manner (**Supplemental Figure 1H**), results that are consistent with established literature^9^. Additionally, OX40 engagement effectively increases T cell production of IL-2 and decreases IL-10, even in the presence of TGFβ (**Supplemental Figure 1I**). Tregs are known to suppress the proliferation of naive T cells *in vitro*^23^. When coculturing Tregs with naïve T cells, we observed that both CD4 and CD8 T cells displayed a significant increase in proliferation upon treatment with αOX40 (**Figure 1I, J**). These data collectively suggest that OX40 agonism has a distinct ability to override suppression through direct effects on conventional T cells themselves, and by negatively regulating the induction of Foxp3 on Tregs.

### OX40 agonism induces a dichotomous response between HNSCC and PDAC during combination radioimmunotherapy

Having established that targeting OX40 generates beneficial effects on T cells *in vitro*, we sought to determine the effect on tumor response in preclinical orthotopic models of HNSCC and PDAC. In MOC2 HNSCC tumors, combination therapy of RT and PD1-IL2v is bolstered by the addition of αOX40, significantly delaying tumor growth compared to RT alone or RT + αOX40 and improving both median and overall survival from 53.5 days to 71 days over RT + PD1-IL2v (**Figure 2A-C**). In contrast, we observed noticeably different effects in orthotopic PK5L1940 PDAC tumors, where the addition of αOX40 to PD1-IL2v negated the therapeutic advantage of combination RT + PD1-IL2v, reducing median survival from 44 days to 29 days (**Figure 2D, E**). The combination of αOX40 with RT also failed to offer benefit to either survival or tumor control, with similar responses seen across models. These results suggest that there are inherent differences between MOC2 and PK5L1940 tumors which ultimately dictate differential responses to our combination radioimmunotherapy.

**Figure 2:**
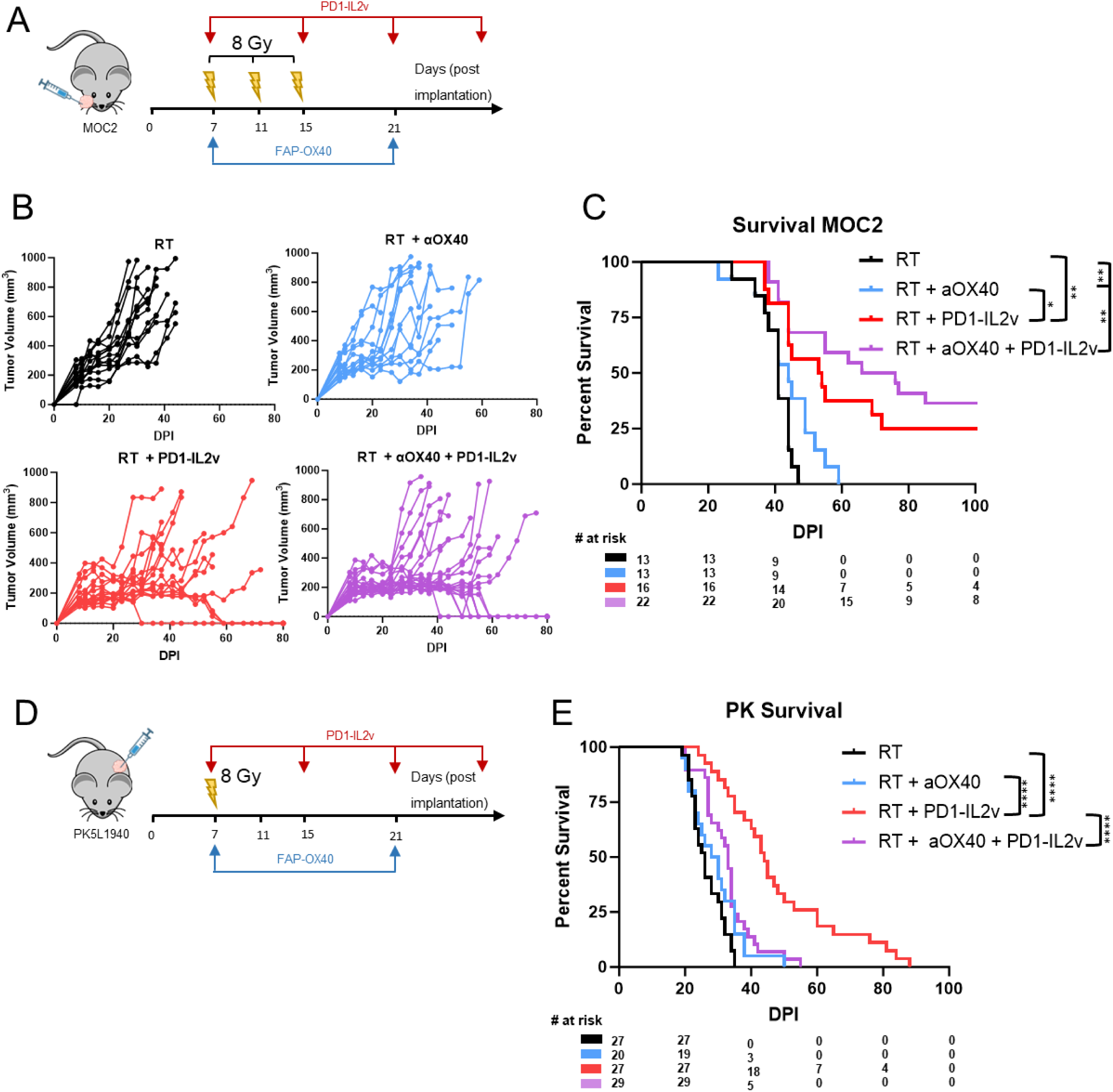
OX40 agonism induces a dichotomous response between HNSCC and PDAC during combination radioimmunotherapy. Experimental Design. C57BL/6 mice were orthotopically implanted with 1×10^5^ MOC2 cells in the buccal mucosa and treated with 3 fractions of 8 Gy RT every 4 days starting at day 7 post implantation. PD1-IL2v was administered via i.p. injection once a week starting at day 7 at a concentration of 0.5mg/kg. αOX40 was administered at day 7 and 21 post implantation at a concentration of 3mg/kg i.p. injection. n=13-22. **B, C)** Tumor growth and survival curves of implanted MOC2 mice during treatment with RT, RT + αOX40, RT + PD1-IL2v, and RT + PD1-IL2v + αOX40. **D)** Experimental Design. C57BL/6 mice were orthotopically implanted with 2×10^5^ PK5L1940 cells in the pancreata and treated with 8 Gy RT at day 7 post implantation. PD1-IL2v was administered via i.p. injection once a week starting at day 7 at a concentration of 0.5mg/kg. αOX40 was administered at day 7 and 21 post implantation at a concentration of 3mg/kg i.p. injection. n=20-27 **E)** Survival curves of mice implanted with PK tumors during treatment with RT, RT + αOX40, RT + PD1-IL2v, and RT + PD1-IL2v + αOX40.

### Accelerated tumor progression is dictated by the tumor microenvironment

Tumors are highly diverse, and while intrinsic cell properties of the cancer cell can drive tumor behavior, for therapies that rely more on immune cell activation to induce cancer cell kill, the makeup of the TME can markedly alter the response to therapy^24,25^. Due to the disparate responses observed during treatment with combination therapy, we wanted to determine how much the discrepancies in the observed response to combination RT with PD1-IL2v plus OX40 agonist are driven by differences in the TME between models.

To interrogate the contribution of the TME to the differential response to combination therapy, we performed heterotopic tumor implantations, where the PDAC cell line was implanted into the HNSCC environment and vice versa. PK5L1940 tumors were implanted into the buccal mucosa of mice and treated with RT and PD1-IL2v or combination therapy (**Figure 3A**). Heterotopic implantation of PK5L1940 tumors resulted in no significant difference in tumor growth between groups that received OX40 agonism compared to those that did not (**Figure 3B**). In contrast, MOC2 HNSCC tumors heterotopically implanted into the pancreas and treated with and without combination therapy yielded the opposite result. A significant decrease in survival was observed paralleling our findings in orthotopic PDAC tumors, with diminished median and overall survival compared to mice treated solely with RT + PD1-IL2v (**Figure 3C, D**).

**Figure 3:**
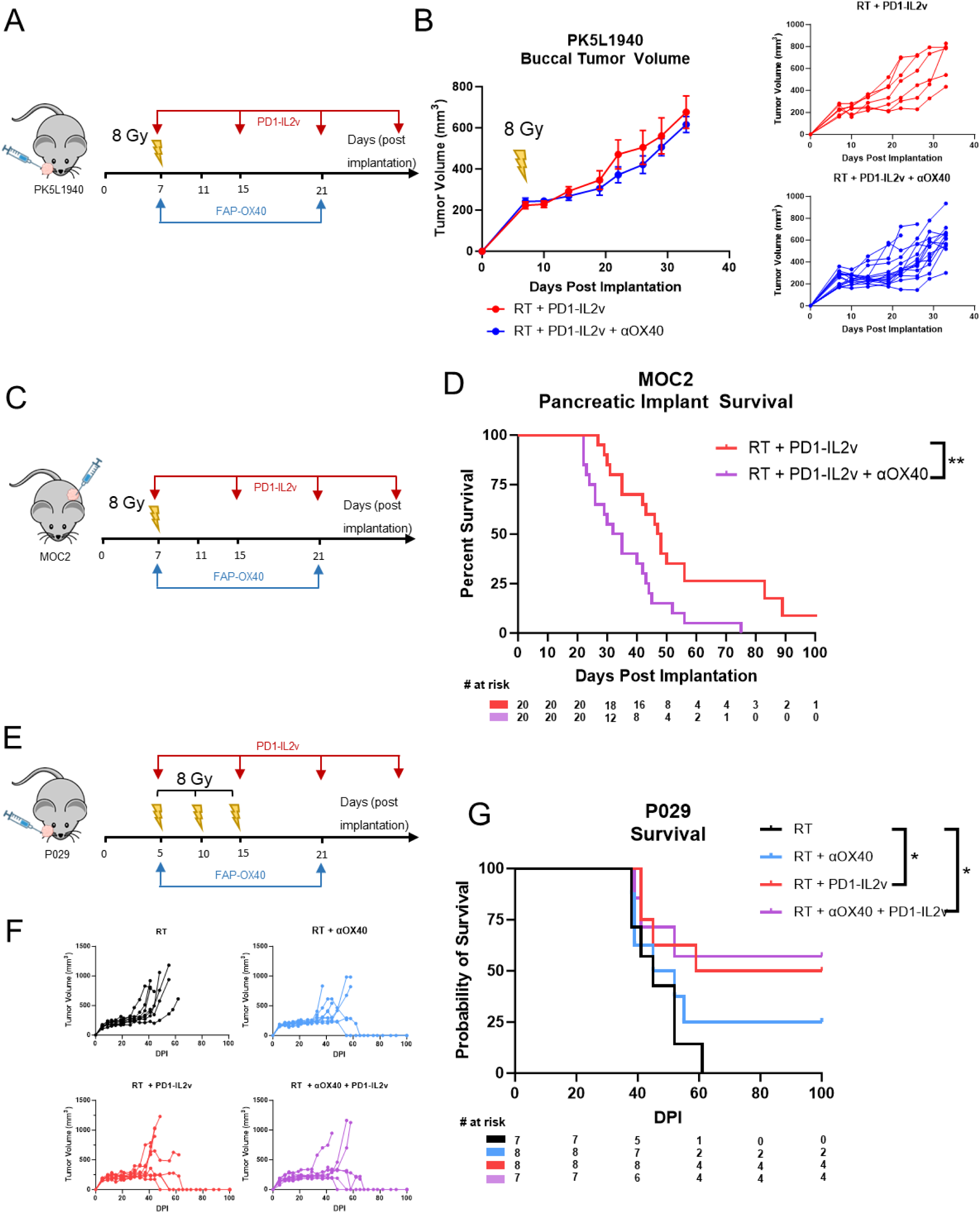
Accelerated tumor progression is dictated by the tumor microenvironment. **A)** Experimental Design. C57BL/6 mice were orthotopically implanted with 2×10^5^ PK5L1940 cells in the buccal mucosa and treated with 8 Gy RT at day 7 post implantation. PD1-IL2v was administered via i.p. injection once a week starting at day 7 at a concentration of 0.5mg/kg. αOX40 was administered at day 7 and 21 post implantation at a concentration of 3mg/kg i.p. injection. n= 7-15. **B)** Tumor growth of implanted PK5L1940 mice during treatment with ,RT + PD1-IL2v, and RT + PD1-IL2v + αOX40. **C)** Experimental Design. C57BL/6 mice were orthotopically implanted with 1×10^5^ MOC2 cells in the pancreata and treated with 8 Gy RT at day 7 post implantation. PD1-IL2v was administered via i.p. injection once a week starting at day 7 at a concentration of 0.5mg/kg. αOX40 was administered at day 7 and 21 post implantation at a concentration of 3mg/kg i.p. injection. n=20. **D)** Survival curves of mice implanted with PK tumors during treatment with RT + PD1-IL2v, RT + PD1-IL2v + OX40, and RT + PD1-IL2v + OX40, +aCD25. **E)** Experimental Design. C57BL/6 mice were orthotopically implanted with 5×10^4^ P029 cells in the buccal mucosa and treated with 3 fractions of 8 Gy RT every 4 days starting at day 7 post implantation. PD1-IL2v was administered via i.p. injection once a week starting at day 7 at a concentration of 0.5mg/kg. αOX40 was administered at day 7 and 21 post implantation at a concentration of 3mg/kg i.p. injection. n= 7-8. **F, G)** Tumor growth and survival curves of implanted mutated P029 mice during treatment with RT, RT + αOX40, RT + PD1-IL2v, and RT + PD1-IL2v + αOX40.

To further control for cancer cell intrinsic genetic mutations as drivers of response to combination immunotherapy, we evaluated the response to combination therapy using the P029 HNSCC line, which shares a similar genetic signature the PK5L1940 PDAC model (**Supplemental Figure 2A**). Kras mutations are the most common genetic perturbations in PDAC, and significant drivers in the development of pancreatic malignancies^26,27^. To rule out tumor progression during combination therapy being a result of cancer cell intrinsic dynamics, and not inherent differences in the TME itself, we orthotopically implanted mice with P029 tumors, that harbor the same Kras^G12D^ mutations as the pancreatic PK5L1940 cell line (**Figure 3E**). Confirming that intrinsic cancer cell properties do not drive response to combination therapy with RT, PD1-Il2v and OX40 agonism, we found that this immunotherapy combination did not elicit the same progression observed in the orthotopic PK5L1940 pancreatic tumors (**Figure 3 F, G**). Rather the response to therapy mirrored that of the head and neck MOC2 cell line (**Figure 2B, C**). Collectively, these data suggest that variations within the TME derived from the site of implantation, and not of cancer cell intrinsic factors, are mediating the response to combination therapy.

### OX40 agonism differentially modulates CD8 T cell responses

Based on the findings that combination immunotherapy elicits opposing effects between tumor models, we aimed to understand the mechanisms that contribute to the diametric responses in orthotopic MOC2 and PK5L1940 tumors. Upon examination of tumor infiltrating lymphocytes, we found that in either tumor model, both CD4 and CD8 T cells had comparable levels of CD44 and IFNγ expression in mice treated with either RT + PD1-IL2v or combination RT + PD1-IL2v and αOX40 (**Figure 4A**). However, the frequency of granzyme B production was significantly dampened in PDAC tumor infiltrating CD8 T cells in mice treated with combination RT+PD1-IL2v and αOX40 compared to PDAC tumors treated with RT+ PD1-IL2v alone or to HNSCC tumors treated with either therapeutic combination (**Figure 4B, C**). Additionally, we see increased expression of the transcription factor TOX, suggesting there may be greater CD8 T cell exhaustion^28^ (**Supplemental Figure 3A**).

**Figure 4:**
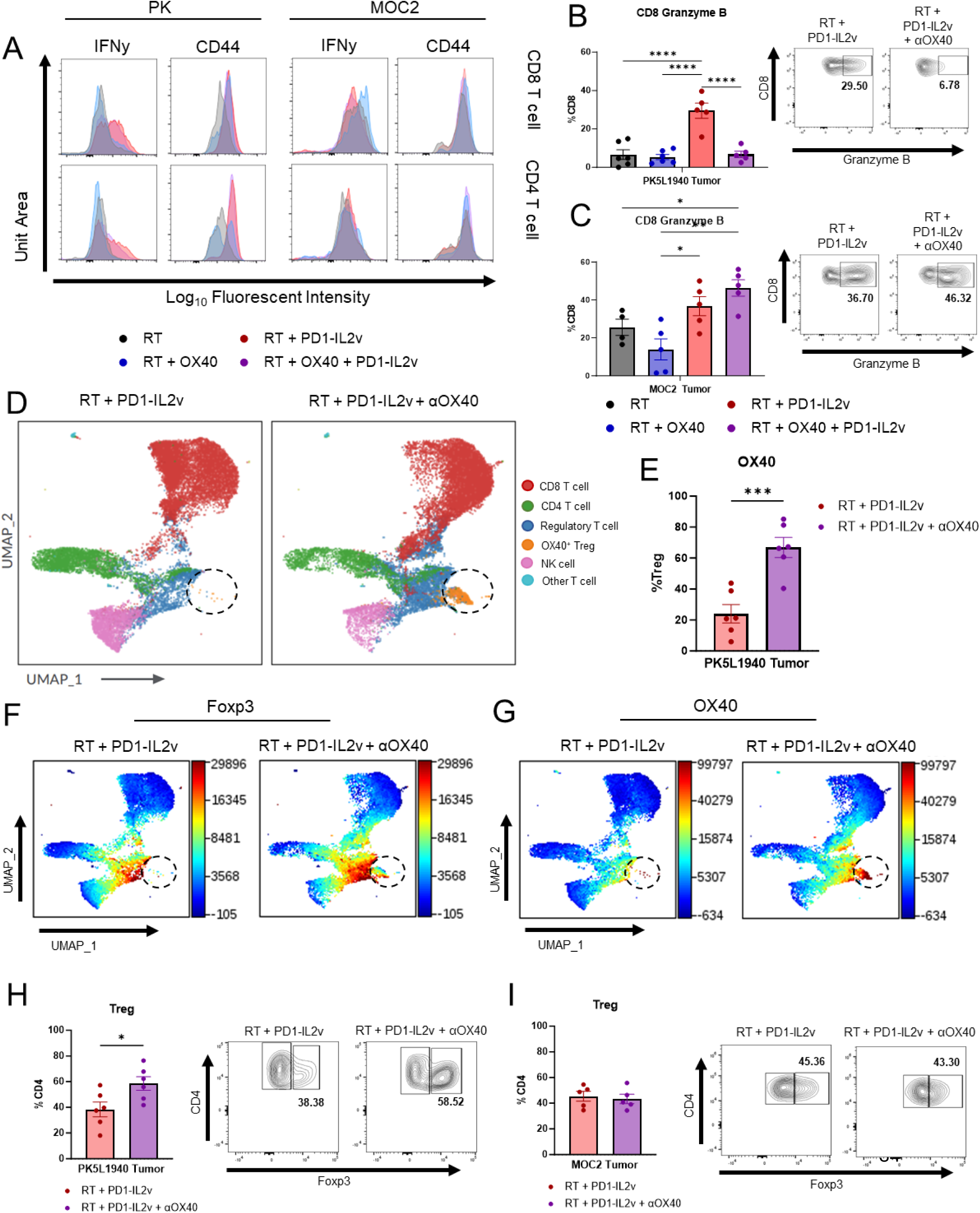
CD8 T cells are differentially modulated and Tregs are enriched in the PDAC TME after combination therapy. **A)** Expression of IFNy and CD44 on intratumoral CD8 (top) and CD4 (bottom) T cells harvested from PK tumor bearing mice harvested 21 past post implantation. **B)** Replicate measurements and representative gating of Granzyme B expression on PK tumor infiltrating CD8 T cells **C)** Replicate measurements and representative gating of Granzyme B expression on MOC2 tumor infiltrating CD8 T cells. **D)** UMAP representation of tumor infiltrating lymphocyte populations from PK5L1940 tumor bearing mice after treatment with RT + PD1-IL2v or RT + PD1-IL2v + OX40. Highlighted is a cluster that is absent in RT + PD1-IL2v only treatment. **E)** OX40 expression on Tregs within the tumor of PK5L1940 tumor bearing mice, harvested 30 days post implantation. **F)** Heatmap representation of Foxp3 expression in PK5L1940 tumor bearing mice between treatment groups. **G)** Heatmap representation of OX40 expression in PK5L1940 tumor bearing mice between treatment groups. **H)** Frequency of regulatory T cells within the tumor of PK5L1950 tumor bearing mice, harvested 30 days post implantation. **I)** Frequency of regulatory T cells within the tumor of MOC2 tumor bearing mice, harvested 21 days post implantation. n=5-6.

As the effects of immunotherapy extend beyond elicited changes within the TME, with systemic immune engagement needed for durable long lasting responses^29–31^, we analyzed the effect that OX40 agonism has on circulating lymphocytes. Unsupervised clustering analysis reveals a significant expansion of circulating CD4 T cells with the addition of αOX40 in MOC2 tumor bearing mice, which was not reciprocated in mice harboring PK5L1940 tumors (**Supplemental Figure 3B, C**). When specifically examining CD8 T cells, we observe comparable levels of CD44 and IFNγ across tumor models. However, granzyme B and Ki-67 expression are notably decreased in MOC2 tumor bearing mice compared to those with PK5L1940 tumors (**Supplemental Figure 3D, E**, hinting that the TME of orthotopic pancreatic tumors is influencing the functional state of CD8 T cells.

### Regulatory T cells are enriched in the PDAC TME after combination therapy

Given the decreased cytotoxicity of PDAC tumor infiltrating CD8 T cells, we sought to investigate the mechanisms underlying the diminished CD8 T cell function. Unsupervised clustering of lymphocytes within the orthotopic PDAC TME reveals a shift in the immune landscape between mice treated with RT and PD1-IL2v and those treated with combination therapy (**Figure 4D**). In mice treated with combination therapy, we identified a T cell cluster exhibiting high co-expression of Foxp3 and OX40, which is notably absent from mice treated with RT and PD1-IL2v alone (**Figure 4 E-G**). Regulatory T cells represent a significant barrier of resistance in both HNSCC and PDAC^18,32–34^, and Treg infiltration into the tumor can potentiate a suppressive TME, increasing tolerance and dampening anti-tumor immunity^18,35–37^. We find a significant enrichment of Tregs within the TME upon treatment with combination therapy in mice harboring PDAC tumors (**Figure 4H**). This increase in Tregs, however, was not replicated in MOC2 tumor bearing mice (**Figure 4I**). These results suggest that, within the PDAC TME, the addition of OX40 directed therapy drives the expansion of a subpopulation of OX40 expressing Tregs, which may contribute to the functionally deficient CD8 T cells observed in the PDAC TME.

### Depletion of regulatory T cells reverses tumor progression in PDAC

Based on the increased prevalence of Tregs within the PDAC TME, and our past work establishing the predilection of Tregs to mediate resistance and tumor progression in our HNSCC and PDAC tumor models^18,33,38^, we hypothesized that Treg depletion would reverse the observed tumor progression during combination therapy (**Figure 5A**). Although it is rarely curative on its own, removal of Tregs from the TME, either through pharmacological depletion or genetic ablation, can effectively lift the restraints on immune mediated tumor cell kill^33,39,40^. Indeed, we observed reversal of the negative effect of OX40 agonism upon the depletion of Tregs via aCD25, completely rescuing the effect of RT + PD1IL2v (**Figure 5B**). Notably, expression of CD25 on Foxp3^+^ cells is significantly reduced, demonstrating effective depletion and penetrance of the antibody into the tumor, a known hinderance to depletion-based therapies (**Figure 5C**).

**Figure 5:**
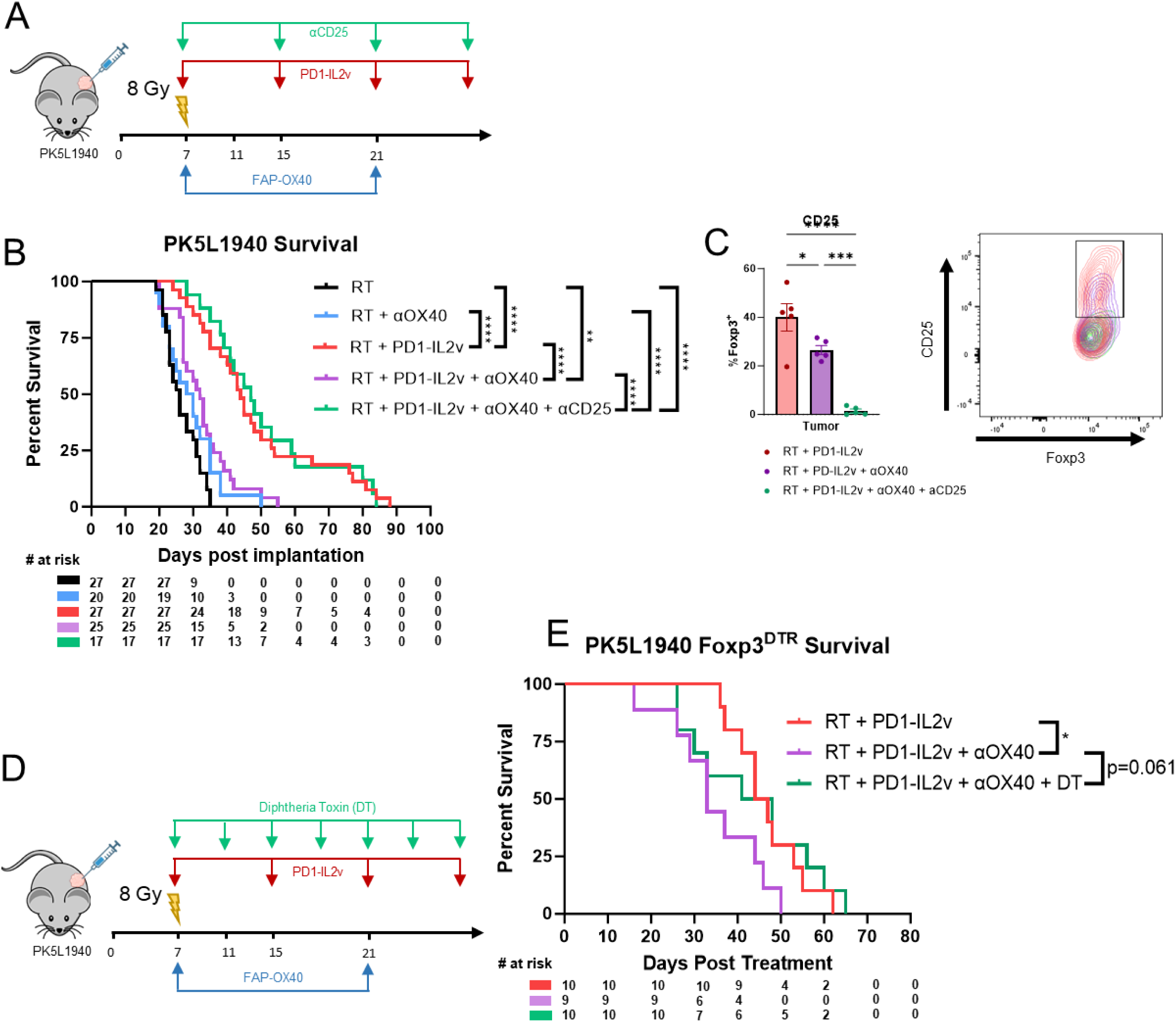
Depletion of regulatory T cells reverses tumor progression in PDAC. Experimental Design. C57BL/6 mice were orthotopically implanted with 2×10^5^ PK5L1940 cells in the pancreata and treated with 8 Gy RT at day 7 post implantation. PD1-IL2v was administered via i.p. injection once a week starting at day 7 at a concentration of 0.5mg/kg. αOX40 was administered at day 7 and 21 post implantation at a concentration of 3mg/kg i.p. injection. αCD25 was delivered via i.p. 1x weekly at a concentration of 3mg/kg. n=17-27. **B)** Survival curves of mice implanted with PK tumors during treatment with RT, RT + αOX40, RT + PD1-IL2v, and RT + PD1-IL2v + αOX40, and RT + PD1-IL2v + αOX40, +αCD25. **C)** Expression and representative flow plots of CD25 on tumor infiltrating Foxp3^+^ Tregs following treatment and depletion using aCD25. **D)** Experimental Design. C57BL/6 or Foxp3^DTR^ mice were orthotopically implanted with 2×10^5^ PK5L1940 cells in the pancreata and treated with 8 Gy RT at day 7 post implantation. PD1-IL2v was administered via i.p. injection once a week starting at day 7 at a concentration of 0.5mg/kg. αOX40 was administered at day 7 and 21 post implantation at a concentration of 3mg/kg i.p. injection. DT was delivered via i.p. 2x weekly at a concentration of 1ug/mouse. n=9-10. **E)** Survival curves of PK5L1940 implanted Foxp3^DTR^ mice treated with RT + PD1-IL2v, RT + PD1-IL2v + αOX40, or RT + PD1-IL2v + αOX40 + DT.

While CD25 is preferentially expressed on Tregs, it is also frequently expressed by effector T cells after TCR stimulation^41^. Therefore, to validate our results and verify that the observed effect is being driven by Tregs and not from off target depletion of conventional CD4 T cells, we utilized the Foxp3^DTR^ mouse model which has a diphtheria toxin receptor (DTR) under the transcriptional control of the Foxp3 promotor. Subsequent administration of diphtheria toxin (DT) results in the selective ablation of Foxp3 expressing cells^42^. Consistent with our previous results, the addition of DT to ablate DTR-expressing Tregs rescued the effect of combination therapy in PK5L1940 tumor bearing mice (**Figure 5D, E**), improving median survival from 32 days to 45.5 days upon Treg depletion.

## Discussion

We provide evidence that the efficacy of orthogonal, combinatorial immunotherapy varies between tumor models and is largely influenced by the composition of the TME. Building upon previous work with models that have significant, although transient, response to combination radiation-immunotherapy, we show that radiation therapy and simultaneous blockade of PD-1 plus stimulation of IL2Rβγ using PD1-IL2v can be successfully combined with OX40 agonism to increase overall tumor control in HNSCC tumors. This effect, however, does not extend to PDAC, where we observed significantly increased tumor progression upon the introduction of an OX40 agonist. Swapping the location of tumor implantation reverses these effects, as does depletion of Tregs, heavily implicating them in dictating response to therapy.

Within the TME, Tregs are widely recognized as a significant barrier of therapeutic response across cancer types, being consistently associated with poorer outcomes^19,43,44^. Directly targeting Tregs has proven to be a challenge as the defining functional feature of Tregs, Foxp3, is localized within the nucleus of the cell, limiting access^45^. Therefore, identifying surface markers which demonstrate pleiotropic effects, disarming Tregs while simultaneously maintaining or enhancing T effector function, is paramount. Unlike the stimulatory action OX40 signaling imparts on conventional T cells, OX40 costimulation on Tregs has classically been associated with diminished suppressive function by inhibiting the induction and expression of Foxp3^9,20,46–48^. These established pleiotropic effects of OX40 signaling on conventional T cells and Tregs, respectively, and our results showing increased OX40 expression on CD8 T cells in both HNSCC and PDAC models informed our decision to target OX40 in the hopes of sustaining the immune activation induced by RT and PD1-IL2v. Our results from orthotopic PK5L1940 model, however, run in opposition to this established dogma and our own included *in vitro* work.

Our findings show that Tregs within the PK5L1940 TME expand upon the addition of OX40 agonism to our backbone treatment of RT and PD1-IL2v. Despite the majority of literature demonstrating that OX40 can effectively reduce Treg suppression, the expansion of Tregs after costimulation with an OX40 agonist was similarly observed by others^48–50^. This may also point to inherent differences in the Treg profile between our HNSCC and PDAC tumor models, as it has been suggested that peripherally induced Tregs (iTregs) are more susceptible to functional destabilization by OX40 simulation that natural thymically derived Tregs (nTregs)^48^. If true, a TME more heavily infiltrated by nTregs rather than iTregs may be more stable and better able to resist disturbances at the Foxp3 locus, potentially accounting for the some of the variance we see across models. Of note, the expansion of Tregs is not present when OX40 agonism is used as a monotherapy and is only seen when used in conjunction with PD1-IL2v. Although the exact mechanism remains to be elucidated, this suggests that the true effect of OX40 simulation on Tregs is highly dependent on context and disease state.

The fundamental role of CD8 T cells in the surveillance and elimination of tumor cells has been well established, with increased infiltration of CD8 T cells being an overwhelmingly positive prognostic marker^51–53^. However, T cells exist across a wide spectrum of phenotypic states, and the presence of CD8 T cells themselves is not indicative of the functional quality of the cell^54,55^. Despite infiltration into the TME, CD8 T cells can become exhausted, a hyporesponsive state characterized by the increased expression of co-inhibitory receptors and progressive loss of effector molecules such as IFNγ and granzyme B^55^. We report a notable reduction of CD8 T cell function in the PDAC TME compared to those infiltrating MOC2 HNSCC tumors. Despite having similar activity in circulation, CD8 T cells infiltrating PK5L1940 PDAC tumors displayed significantly reduced granzyme B expression and TOX expression, a key transcriptional regulator of exhaustion^28^, upon the introduction of OX40 agonism to our therapy. These data seem to suggest that the induction of CD8 T cell exhaustion and attenuation of tumor reactive CD8 T cell function is a possible consequence of the increased Treg abundance in the PDAC TME in the setting of our combination therapy.

Overall, the outcome of this work presents two substantial findings. First, we show that despite both preclinical HNSCC and PDAC displaying significant therapeutic benefit from RT and PD1-IL2v^14,15^, their response upon the addition of αOX40 act in opposition. MOC2 HNSCC tumors receive additional benefit from combination therapy, driving further tumor eradication. PK5L1940 tumors, in contrast, exhibit a diametric response compared to MOC2 with significantly reduced survival compared to RT and PD1-IL2v. This effect we show is due to increased prevalence of intratumoral Tregs, as depletion abrogates tumor progression. Next, and perhaps most importantly, we demonstrate through heterotopic swapping of tumor implantations that the location of the tumor heavily influences overall response to combinatorial radio-immunotherapy.

As more interventions become available in the oncologic armamentarium, combination immunotherapies are becoming more commonplace. Our data demonstrate that combination immunotherapies can assert unexpected negative consequences depending on the context of tumor type, implantation site, and overall composition of the TME.

## Declarations

### Ethics approval

All protocols for in vivo animal studies were approved by the Institutional Animal Care and Use Committee (IACUC) of the University of Colorado Anschutz Medical Campus

### Data Availability

The data generated in this study are available upon request from the corresponding author.

### Author Contributions

**J. Gadwa:** Conceptualization, data curation, formal analysis, methodology, visualization, writing-original draft. **J. Yu:** Formal analysis, investigation, methodology, writing-review and editing. **M. Piper:** Formal analysis, investigation, methodology, writing-review and editing**. M.W. Knitz:** Data curation, methodology, software. **M. Pincha:** Formal analysis, investigation, methodology. **C. Klein:** Visualization, resources, methodology, writing-review and editing. **M. Amann:** Visualization, resources, methodology, writing-review and editing. **S.D. Karam:** Conceptualization, supervision, funding acquisition, visualization, writing-review and editing, writing-original draft.

## Supplemental Figures

**Supplemental Figure 1:**
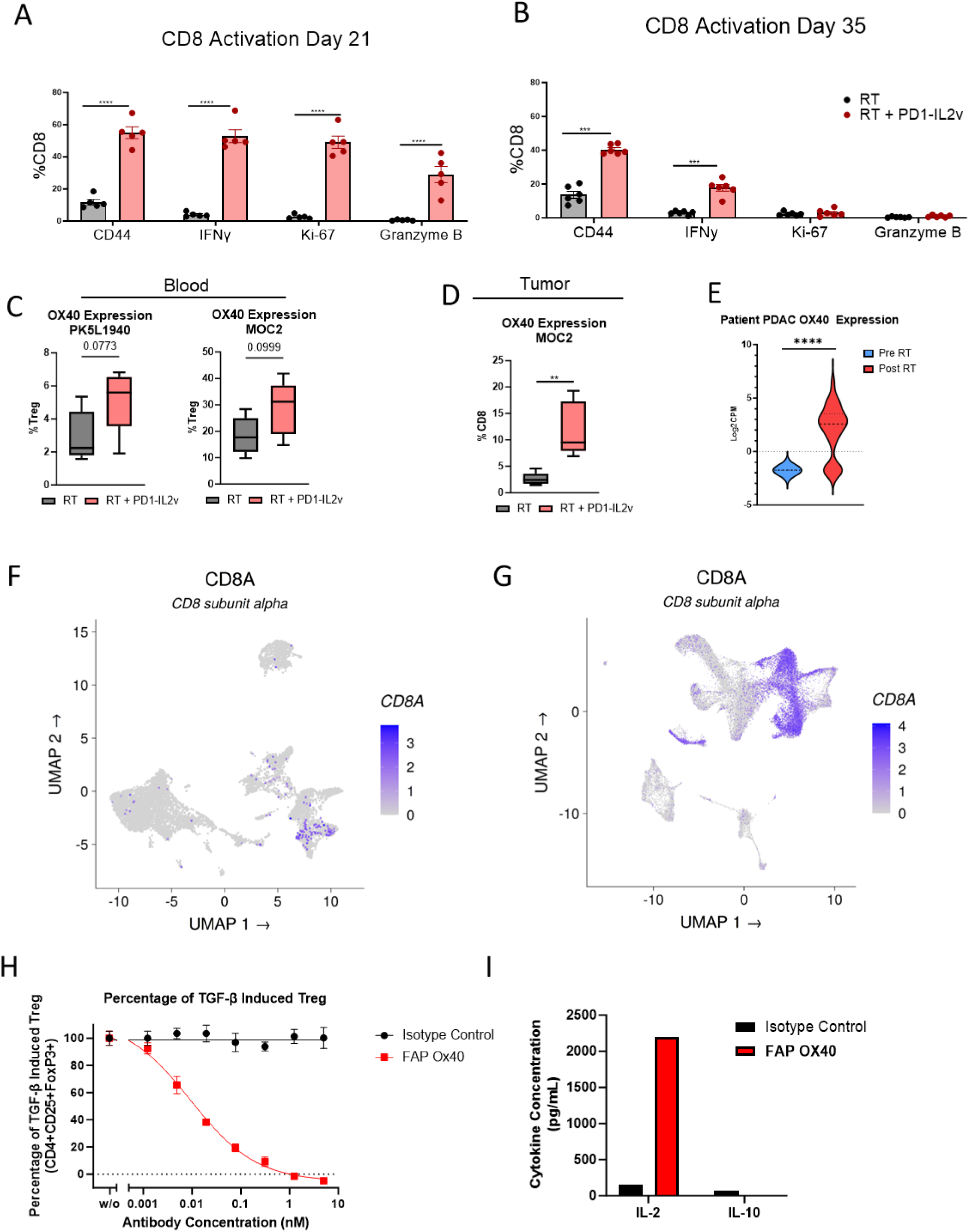
**A)** Expression of CD44, IFNγ, KI-67, and Granzyme B from circulating CD8 T cells 21 days post implantation. **B)** Expression of CD44, IFNγ, KI-67, and Granzyme B from circulating CD8 T cells 35 days post implantation **C)** Expression of OX40 on circulating Tregs in PK5L1940 and MOC2 tumor bearing mice **D)** Expression of OX40 on tumor infiltrating CD8 T cells from MOC2 tumor bearing mice. **E)** Expression of OX40 among clinical PDAC and HNSCC patients before and after treatment with radiation therapy **F)** UMAP analysis of sn-RNA sequencing showing CD8a expression on immune cell clusters from patients with pancreatic ductal adenocarcinoma **G)** UMAP analysis of sc-RNA sequencing showing CD8a expression on immune cell clusters from patients with head and neck squamous cell carcinoma. **H)** Percentage of Treg induction after treatment with TGF-β and either isotype control or αOX40.

**Supplemental Figure 2:**
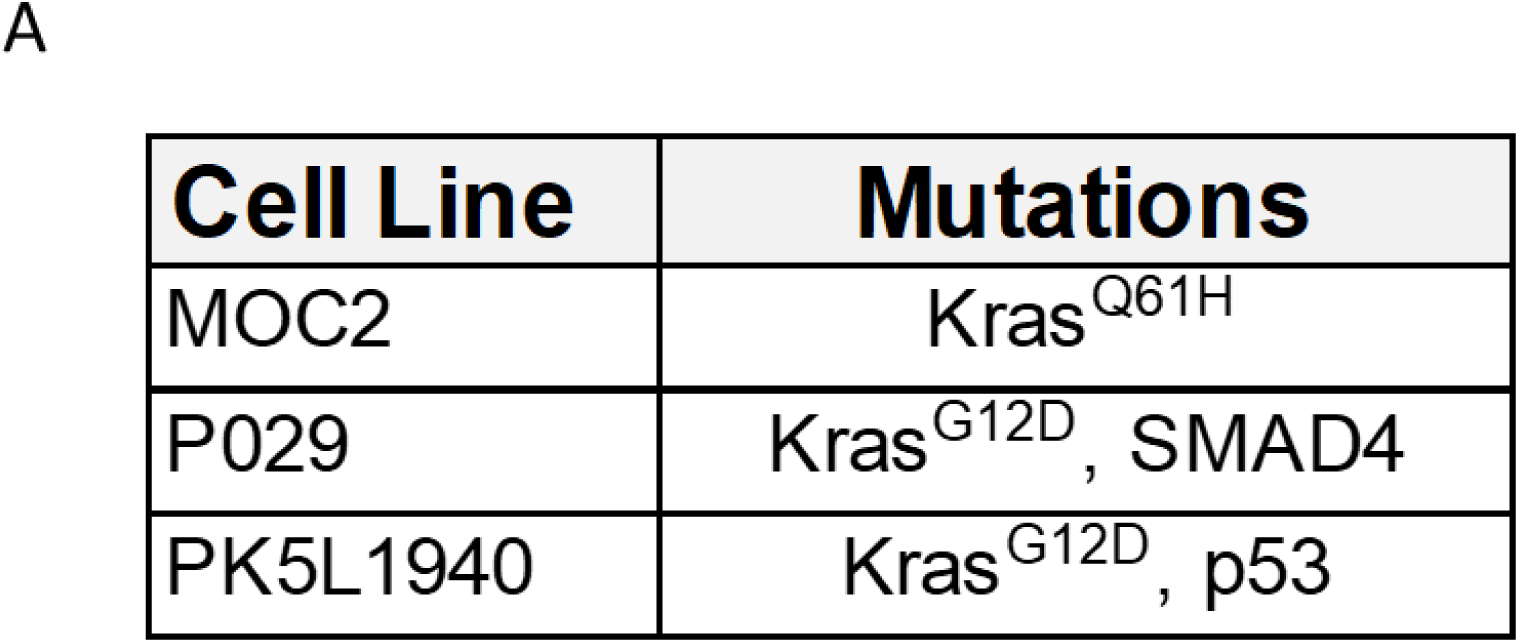
**A)** Table showing the genetic mutations of MOC2, P029, and PK5L1940 cancer cell lines

**Supplemental Figure 3:**
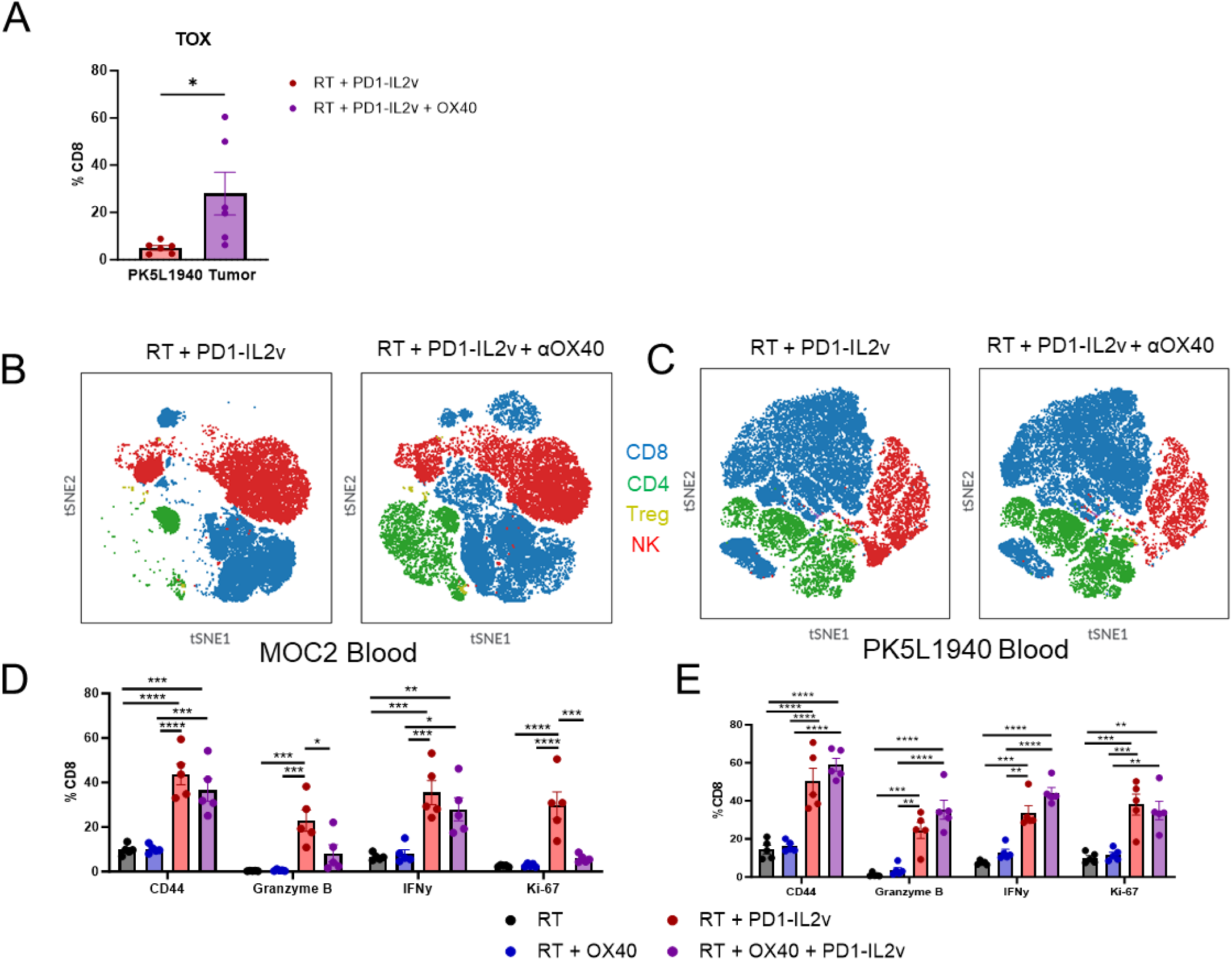
**A)** Expression of the transcription factor TOX in CD8 T cells from tumor bearing PK5L1940 mice. **B)** tSNE representation of circulating T cells and NK cells from MOC2 tumor bearing mice. **C)** tSNE representation of circulating T cell and NK cells from PK5L1940 tumor bearing mice. **D)** Expression of CD44, Granzyme B, IFNy, and Ki-67 on circulating CD8 T cells from MOC2 tumor bearing mice. **E)** Expression of CD44, Granzyme B, IFNy, and Ki-67 on circulating CD8 T cells from PK5L1940 tumor bearing mice.

